# Modeling ex vivo tumor-infiltrating lymphocyte expansion from established solid malignancies

**DOI:** 10.1101/2021.02.04.429846

**Authors:** HM Knochelmann, AM Rivera-Reyes, MM Wyatt, AS Smith, R Chamness, CJ Dwyer, M Bobian, GO Rangel Rivera, JD Horton, M Lilly, MP Rubinstein, DM Neskey, CM Paulos

## Abstract

Adoptive transfer of tumor-infiltrating lymphocytes (TIL) elicits the regression of metastatic malignancies, yet a low proportion of patients achieve complete durable responses. The high incidence of relapse in these patients highlights the need to better understand mechanisms of tumor escape from T cell control. While melanoma has provided the foundation for developing TIL therapy, much less is known about TIL efficacy and relapse in other malignancies. We sought to investigate TIL characteristics in mouse tumors which have not been studied in this setting. Here, we expanded murine TIL *ex vivo* in IL-2 from fragments of multiple tumor models, including oral cavity cancer models of varying immunogenicity. Additionally, TIL was expanded from pmel-1 mice bearing B16 melanoma, yielding an enriched population of tumor-infiltrating TCR transgenic T cells. Murine TILs are similar to human TIL in that they express high levels of inhibitory receptors (PD-1, Tim-3, etc.) and can be expanded *ex vivo* in IL-2 extensively. Of clinical relevance, we draw parallels between murine and patient TIL, evaluating relationships between PD-1, Lag-3, and Tim-3 on TILs from a cohort of oral cavity cancer patients. This platform can be used by labs even in the absence of clinical specimens or clean cell facilities and will be important to more broadly understand TIL phenotypes across many different malignancies.

## Introduction

The adoptive transfer of large numbers of tumor-infiltrating lymphocytes (TIL) is a promising clinical immunotherapy strategy as objective responses of up to 50% have been reported in patients with metastatic disease (1, 2). Modern TIL expansion protocols were derived from early discoveries that large numbers of T cells could mediate long-lived immunity to tumors, especially when supported by exogenous IL-2 (3, 4). Clinically, TILs are expanded *ex vivo* in IL-2 for several weeks, followed by a rapid expansion period where cells can expand up to 1000-fold (5). While advancements in TIL culture and selection for tumor-reactivity have supported efficacy in some patients, the long-term duration of T cell culture has made translation difficult, thus highlighting the continued need for optimized processes related to T cell product development.

Multiple methods of TIL expansion have been described, each seeking to improve cell yield and reactivity to tumors. Two distinct methods include either culturing single-cellular digests of TILs and tumor cells, or generating “tumor microcultures;” the latter allows T cells to egress from tumor fragments, and is more efficient for identifying tumor-reactive clones (5). While the cell-suspension method has been routinely tested in murine sarcoma, leukemia, and colorectal cancer models since the 1980s (3, 4), the feasibility of establishing tumor microcultures in other murine cancers as preclinical models of TIL therapy is relatively undescribed.

Beyond direct isolation of T cells from mouse tumors, TIL models have expanded to include adoptive transfer of transgenic T cells, where T cells in mice express T cell receptors (TCRs) specific to antigens expressed in tumor lines (6-9). Two transgenic models for melanoma, the pmel-1 and the TRP-1 models, were created to provide a source of T cells which target naturally expressed melanocyte differentiation antigens (gp100 or tyrosinase proteins) (8, 9). Other novel methods of generating T cell-transgenic mice include nuclear transfer models, where mice are designed to express specific TCRs rearranged naturally against antigens of interest (10, 11). These models have been a critical proxy for TIL therapy as they allow study of tumor-reactive T cells, their function, and novel methods to enhance immunity against aggressive cancers.

However, there remain disadvantages for using TCR transgenic mice to model TIL therapy. Transgenic T cells are often obtained from the animal’s spleen or lymph nodes, which provides a source of antigen-specific T cells, yet the cells have not encountered tumor unlike a patient’s TIL. As T cells obtained peripherally are likely to be phenotypically distinct from those that reside in the tumor, the starting T cell product may be less clinically relevant compared to patient TIL. Additionally, TIL models for cancers beyond melanoma are underdeveloped; therefore, there remains poor understanding of the mechanisms underlying regression and relapse after ACT therapy in other solid tumors.

Herein, we sought to generate both natural and transgenic murine TIL in a clinically relevant manner to study characteristics of TIL which may influence their ability to elicit antitumor responses. We expanded TIL from multiple mouse tumors by culturing small tumor fragments *ex vivo* in IL-2. TIL expansion efficiency using this method differed across tumor origin, as did the composition of immune cells expanding from the tumor. TILs were also successfully expanded *ex vivo* from tumors on transgenic pmel-1 mice, bearing B16F10 tumor against which their TCRs are reactive. Phenotypically, TILs expanded from tumors have higher expression of inhibitory receptors (IRs) like PD-1 and Tim-3 (12) than T cells isolated from peripheral tissues. Finally, we draw parallels from murine TIL to patient TIL, using a cohort of patients with oral cavity squamous cell carcinoma (OCSCC), and describe the inhibitory receptor expression profiles of CD4^+^ and CD8^+^ TIL products from this population. Overall, using murine tumor fragments to expand TIL provides a platform to evaluate therapeutic outcomes in combination with checkpoint blockade or other immune modulating strategies in weakly immunogenic tumors.

## Materials and Methods

### Animal studies

C57BL/6 mice were purchased from Jackson labs or Taconic labs, and pmel-1 TCR transgenic mice (B6.Cg-*Thy1*^*a*^/Cy Tg(TcraTcrb)8Rest/J) were purchased from Jackson labs. Male or female mice were bred and housed in the Hollings Cancer Center at the Medical University of South Carolina. Housing and experiments were conducted in accordance with MUSC’s IACUC procedures and with the supervision and support of the Division of Laboratory Animal Resources.

### Tumor lines

Moc2 and Moc22 cell lines were obtained from Ravindra Uppaluri at the Dana-Farber Cancer Institute and were cultured in sterile media containing the following elements as described previously (13): 2:1 mixture of IMDM: Hams nutrient mixture containing 1% Pen/Strep, 5% FBS, and 5ug/mL insulin, 5ng/mL EGF, and 40ng/mL hydrocortisone. LLCA9F1 was a gift from Mark Rubinstein and Eric Bartee at the Medical University of South Carolina and was cultured with DMEM containing 10% FBS and 1% Pen/Strep. B16F10 (haplotype H2b) was a gift from Nicholas P. Restifo, NCI Surgery Branch, and were cultured in RPMI complete medium. B16F10 and Moc cell lines were confirmed mycoplasma and pathogen free most recently in March 2020.

### Tumor transplantation

All cell lines were injected subcutaneously in PBS in the abdomen of recipient animals. Moc22 and Moc2 were injected into Taconic mice and dosed at 1×10^6^ and 0.1×10^6^ cells per mouse respectively due to differences in growth kinetics. LLC was given to C57BL/6 from Jackson labs in a dose of 0.4-0.5×10^6^ cells/mouse. B16F10 was given in doses of 0.4-0.5×10^6^ cells/pmel mouse.

### Ex vivo TIL expansion

Murine tumors were harvested prior to tumor size reaching 10×10mm^2^. Tumors were removed from mice under sterile conditions and incubated in sterile RPMI complete medium on ice. Mouse tumors were then diced into pieces measuring 3-4mm^2^, washed, and plated in media containing high dose IL-2 (6000IU/mL) prior to incubation at 37°C. Cultures were left undisturbed for a minimum of 5 days, after which half of the media was changed at least 2-3 times per week. TILs were split upon reaching confluency and were maintained at approximately 1×10^6^ cells/mL. The remaining tumor was removed at the time of first split or at three weeks into culture, depending on rate of cell egress from the tumor. All TIL cultures were maintained as individual cultures after plating from a distinct tumor piece. Human TIL was generated in a similar manner, modeling off protocols established by others previously (5).

### Tissue distribution

Tissues were collected from tumor-bearing animals to identify characteristics of T cells existing peripherally compared to those expanded from the tumor microenvironment. Peripheral blood was collected via small lancet puncture of the mandibular vein into a tube containing 0.125 M EDTA. Spleens were collected into RPMI complete medium, processed to a single cell suspension over a 70uM filter. Remaining RBCs were lysed, and samples were filtered prior to analysis.

### Clinical Samples

Samples were collected after informed consent and were deidentified prior to delivery to the research lab. All studies were approved by the Institutional Review Board at the Medical University of South Carolina prior to initiation.

### Flow Cytometry

BDFACSVerse instruments were used for flow cytometry data collection, and analysis was conducted using FlowJo software (BD). Samples were stained for extracellular proteins by suspension in FACS buffer (PBS + 2% FBS) with Fc block (1:500 dilution), and incubation with antibodies at 4°C for 25 minutes. For cytokine production analysis, samples were activated with aCD3 agonist (clone 145-2C11, BioLegend) overnight and incubated with Monensin (2μM) and Brefeldin A (5μg/mL) (Biolegend) for 4 hours, followed by fixation and permeabilization according to manufacturer’s protocol (BioLegend). Unstimulated controls were included for all assays. A complete list of antibodies is found in the **Supplementary Materials**.

### Statistical Analysis

Statistical analysis was performed with Graphpad Prism (v7.0). Continuous measures compared between two groups were made using Mann Whitney U test as assumptions for the t-test were most often not met. For functional comparisons of T cell subsets expressing inhibitory receptors, the Wilcoxin signed-rank test was used as a pairwise comparison within biological replicates. Given the data are exploratory in nature and it was desired not to be overly restrictive, hypothesis tests were not adjusted for multiple comparisons. P values less than 0.05 were considered significant. Plots display mean values and error bars represent standard deviations. Outliers were not omitted in the data presented.

## Results

### Modeling clinical TIL expansion with murine solid tumors

Human T cells egress from tumor fragments when placed in high doses of IL-2; therefore, we posited that murine TILs could be expanded from tumors in a similar manner. We implanted three different tumor cell lines on syngeneic immunocompetent mice, including two oral cavity cancer lines (Moc2 and Moc22) and one Lewis Lung carcinoma cell line (LLCA9F1). After growing for at least one week on mice (**Fig. 1A**), tumors were harvested and processed in a sterile manner by dicing the tumor into small fragments approximately 3-4mm^2^. Each tumor fragment was rinsed in fresh media and then plated in individual wells of a 24-well plate in 2mL of media with 6000IU IL-2/mL (**Fig. 1B**). T cells were allowed to egress from the tumor and were expanded in culture as described in *Methods*.

**Figure 1:**
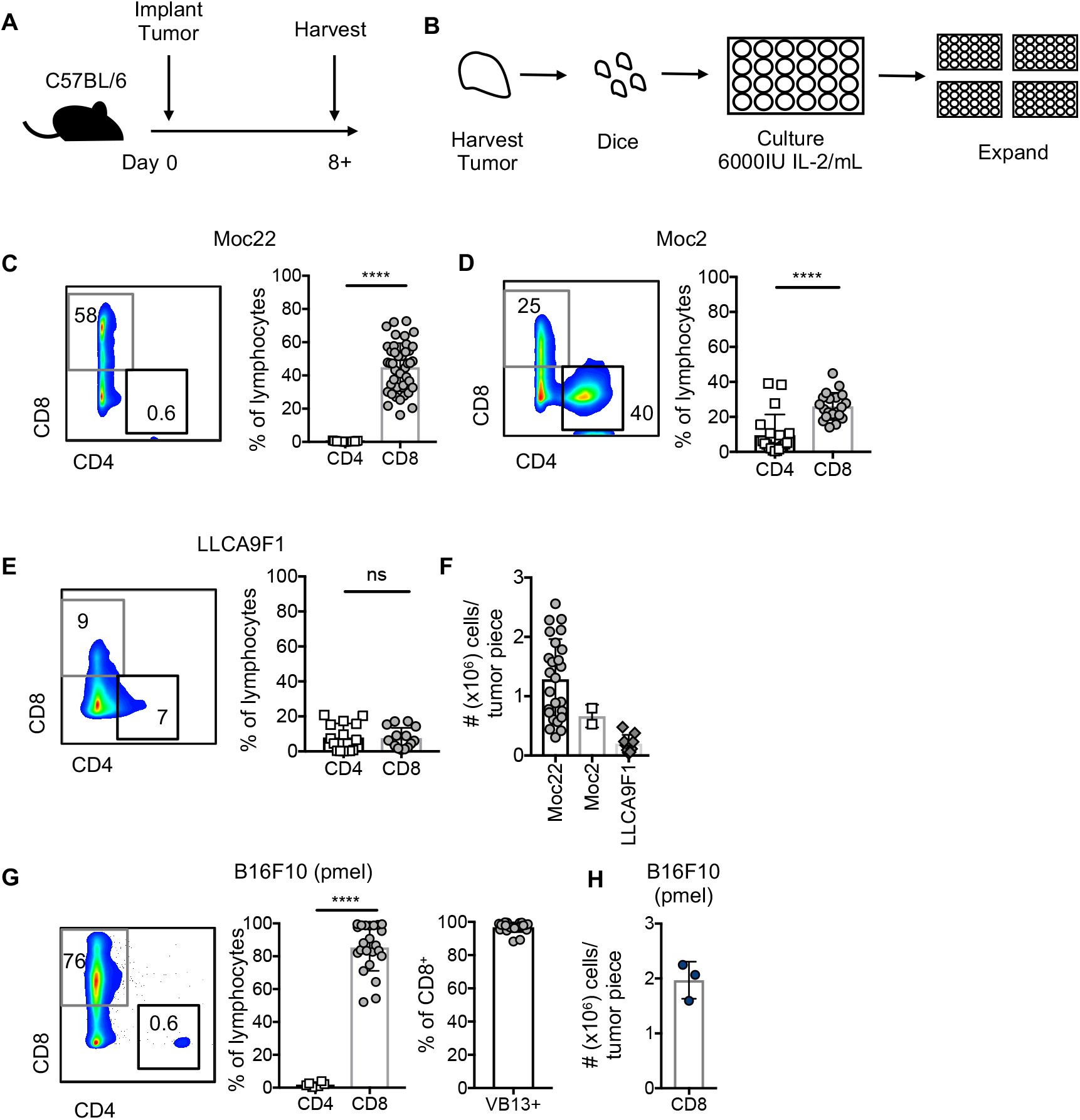
Characteristics of TIL expanded *ex vivo* from murine solid tumors. A) Model schematic. Tumors are established in mice for one week or more prior to harvest for TIL expansion. B) Schematic of ex vivo TIL expansion. Tumors are diced into ∼3-4mm^2^ fragments. Each fragment is plated in a separate well of a 24-well plate with media containing 6000IU IL-2/mL. These cells are expanded 3 weeks prior to analysis. C) T cell populations expanded from Moc22 tumors. n=43 TIL cultures from 26 mice. D) TIL populations expanded from Moc2 tumors. n=20 TIL cultures from 10 mice. E) T cell populations expanded from LLCA9F1 tumors; n=15 cultures from 5 mice. F) Yield of cells after 3 weeks of expansion based on tumor cell line. G-H) TIL expanded from B16F10 tumors established on pmel-1 transgenic mice, where Vβ13 marks the TCR specific for melanoma. n=23 TIL cultures from 3 mice. Statistical tests: C-F, H, Mann Whitney U test; ****p<0.0001; ns, not significant.

After three weeks of expansion, TILs were collected and assayed for T cell phenotypes. Overall, we found that the frequencies of CD4^+^ and CD8^+^ T cells were variable and dependent upon tumor origin. Within the oral cavity cancer cell lines, TIL expanded from Moc22 tumors was predominantly CD8^+^ T cells, while Moc2 tumors had a mix of CD4^+^ and CD8^+^ T cells (**Fig. 1C-D**). These findings are concordant with previously reported T cell populations infiltrating OCSCC tumors (14). In contrast, TILs expanded much less efficiently from LLCA9F1 using this protocol (**Fig. 1E**). Total yields of CD8^+^ T cells were higher for oral cavity cancers relative to lung cancers and ranged from <0.1-3×10^6^ cells per 3-4mm^2^ tumor fragment (**Fig. 1F**).

As the OCSCC and LLC TILs were derived from C57BL/6 hosts, we next questioned whether transgenic, antigen-specific CD8^+^ T cells could be expanded from a tumor growing on a pmel-1 TCR transgenic mouse. Previous studies have demonstrated that a B16F10 melanoma tumor grows unabated on the pmel mouse(8), yet the ability to expand TIL *ex vivo* from this mouse remains unexplored. To address this, B16F10 tumors were established on pmel-1 transgenic mice, where the CD8^+^ T cells within the animal express an H2-D^b^ restricted TCR specific for gp100, expressed in melanoma and healthy melanocytes (8). Indeed, expansion of pmel-T cells from B16F10 tumors was efficient, yielding predominantly CD8^+^ T cells, nearly all of which expressed Vβ13 of the gp100-specific TCR (**Fig. 1G**). Total yield of pmel T cells neared approximately 2×10^6^ cells per tumor fragment (**Fig. 1H**). Thus, TILs can be expanded *ex vivo* from murine tumors using protocols mirroring clinical TIL expansion, and TIL composition varies across tumor cell lines, likely related to the immunogenicity of the malignancy itself.

### Murine TIL are enriched in inhibitory receptors relative to peripheral T cells

We next sought to determine whether TIL had different inhibitory receptor expression profiles relative to peripheral T cells. To address this, we focused on Moc22 and Moc2 cell lines as TIL were reproducibly expanded from these tumors. As shown in the schematic in **Figure 2A**, Moc2 or Moc22 tumors were established for 12 days, after which tumors and peripheral blood were collected. TIL were expanded *ex vivo* from tumors for 4 weeks and assayed for surface marker expression compared to peripheral blood (**Fig. 2A**). As expected, CD8^+^ TILs expressed a significantly higher Tim-3 and PD-1 compared to peripheral blood from the host of origin (**Fig. 2B-C**). Interestingly, in Moc22 tumors, up to 40% of TILs co-expressed these inhibitory receptors while in Moc2 tumors, less than 20% co-expressed PD-1 and Tim-3 (**Fig. 2B-C**). While expression of these markers in tumor-infiltrating T cells has been reported previously (15), it remained possible that the high doses of IL-2 could have driven some of the differences in phenotype we observed between TIL and peripheral blood.

**Figure 2:**
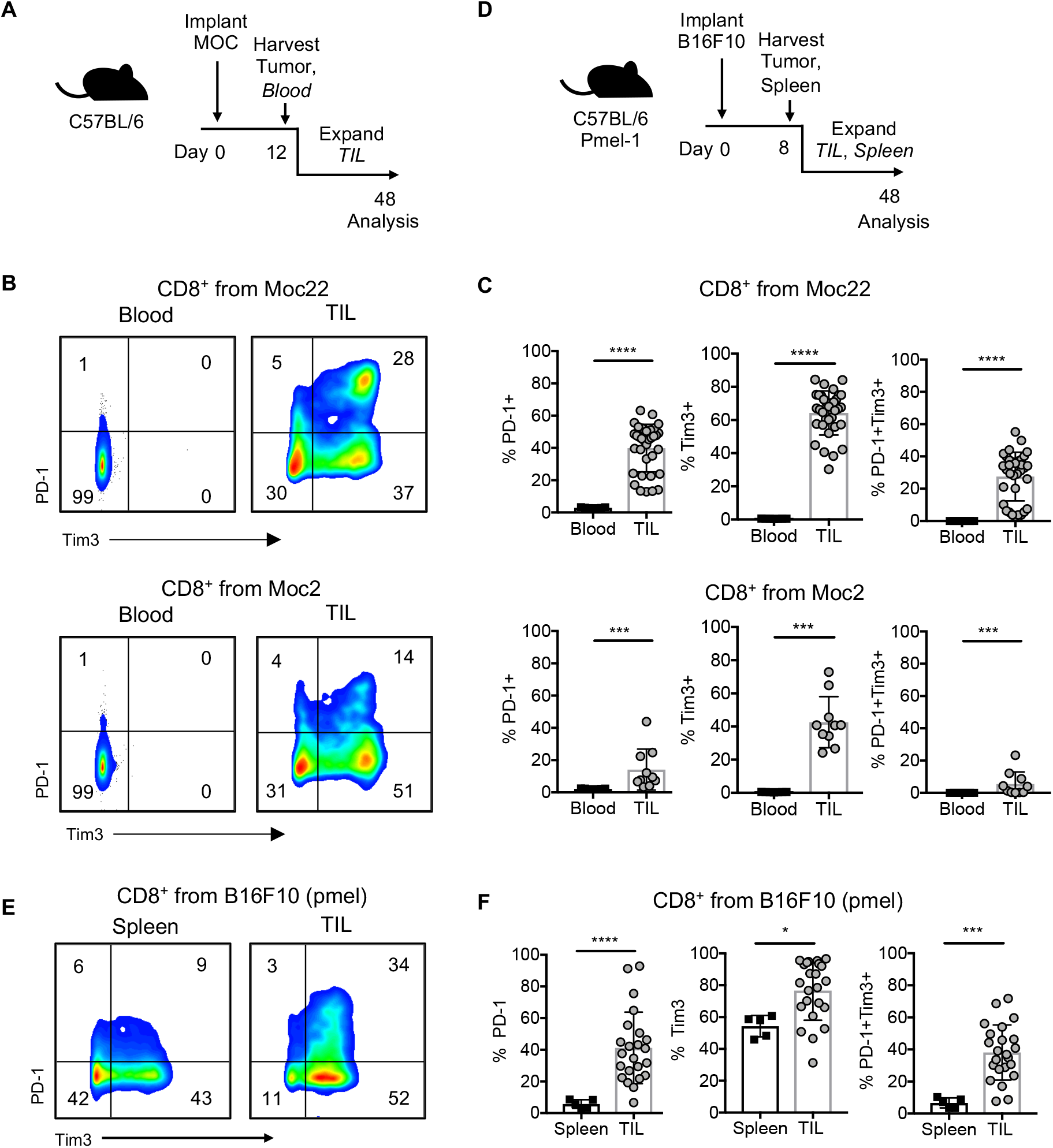
Inhibitory receptor expression in murine TIL exceeds peripheral T cells. A) Moc2 and Moc22 tumors were established to a palpable subcutaneous tumor in C57BL/6 animals for 12 days prior to tumor harvest and plating for TIL expansion. Blood was obtained from time of harvest for a peripheral T cell control. B-C) Inhibitory receptor expression in Moc22 and Moc2 CD8^+^ TIL relative to blood. Blood: n=7, TIL: Moc22 n=34 cultures from 21 mice, combined from two independent experiments, Moc2 n=10 cultures from 5 mice. D) Pmel-1 transgenic mice were inoculated with B16F10 melanoma tumor. Transgenic T cells infiltrating tumors or spleens were expanded ex vivo and characterized. E-F) Surface expression of inhibitory receptors after 3-4 weeks of expansion in IL-2. Spleen n=5, TIL cultures n=23 from 3 mice. Statistical tests: C,F) Mann Whitney U test, *p<0.01, ***p<0.001, ****p<0.0001.

To account for the potential impact of IL-2 on T cell phenotypes, we expanded TIL from either the tumor or spleen of the pmel transgenic animal in high doses of IL-2 (**Fig. 2D**). We found, after 4 weeks of expansion in IL-2, approximately 80% of pmel TILs expressed Tim-3 while approximately 40% expressed PD-1 or co-expressed the two (**Fig. 2E-F**). While spleen-derived T cells, expanded in high dose IL-2, expressed Tim-3 and PD-1, they still did not express these inhibitory receptors to the extent of T cells expanded from the tumor (**Fig. 2E-F**). Thus, tumor exposure was critical to drive high PD-1 and Tim-3 expression on TILs.

### Cytokine production is highest with ex vivo stimulation in TILs co-expressing PD-1 and Tim-3

We next questioned whether cytokine production in *ex vivo* expanded TILs differed in CD8^+^ T cell subsets which either singly expressed, co-expressed, or expressed neither PD-1 nor Tim-3 (**Fig. 3A**). We hypothesized that T cells co-expressing these markers represented exhausted T cells, which would have the lowest effector function upon TCR stimulation(16). To address this, TILs expanded from Moc22 tumors were activated with plate bound aCD3 agonist overnight. As a marker of activation, we examined forward and side scatter of the IR subsets. As expected, PD-1^+^Tim-3^+^ double positive T cells had the largest cell size and highest granularity, while cells negative for both markers had the smallest size and granularity (**Fig. 3B**).

**Figure 3:**
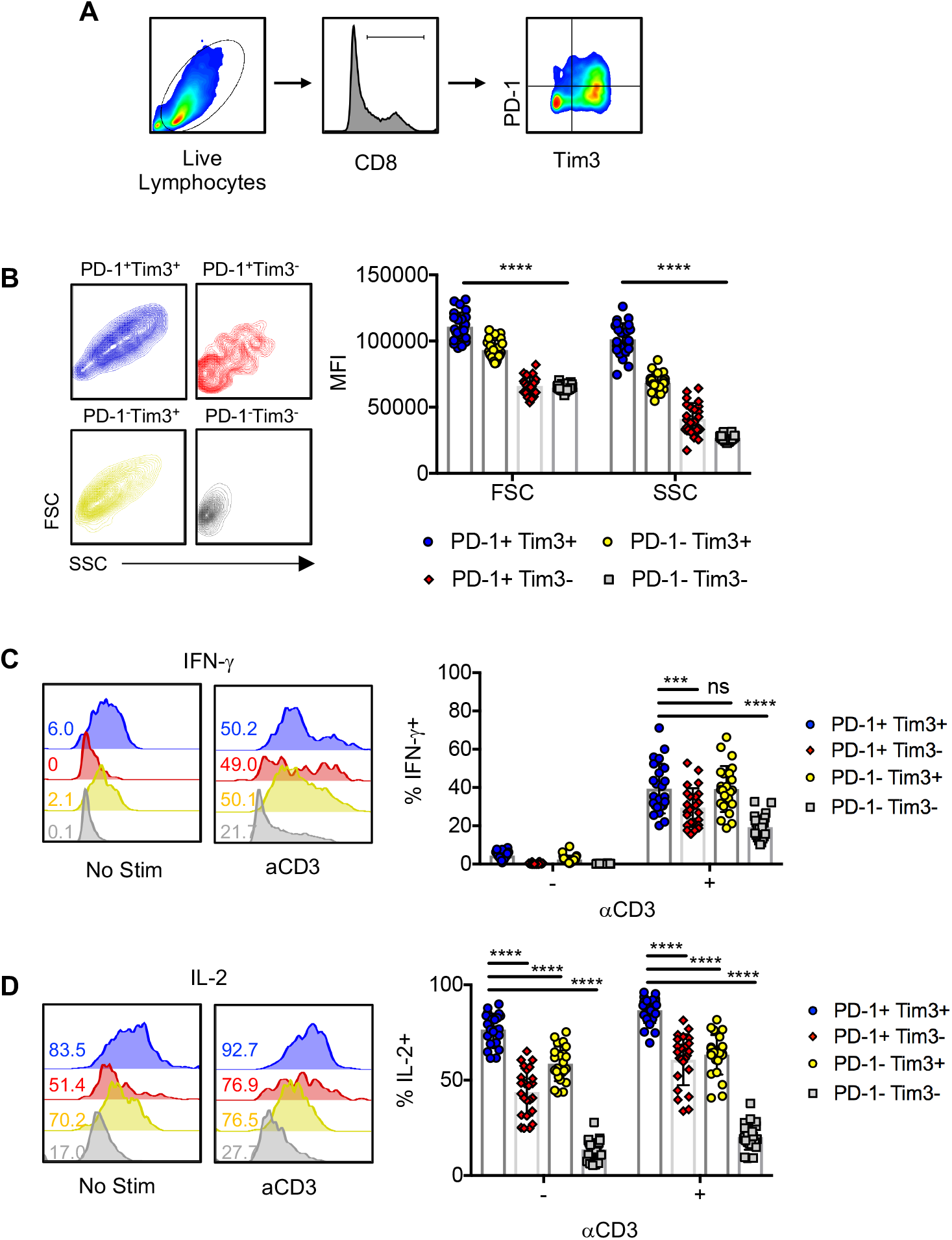
*In vitro* cytokine production is highest in TIL expressing PD-1 and Tim-3. A) Gating strategy. Lymphocytes expanded from Moc22 tumors were gated on live cells, FSC/SSC and CD8 expression prior to gating for expression of Tim-3 or PD-1. Cells were activated with 1mg/mL plate-bound aCD3 overnight with Golgi stop prior to analysis by flow cytometry. B) FSC and SSC of cell populations expressing PD-1, Tim-3, or co-expressing these markers. C-D) Histogram and biological replicates displaying C) IFN-γ or D) IL-2 production from cells activated overnight or at resting state by inhibitory receptor expression. n=24 TIL cultures from 16 mice. Statistical tests: Wilcoxon matched-pairs sign rank test, ***p<0.001, ****p<0.0001.

Next, we examined cytokine production in these subsets. After determining frequency of IFN-γ and IL-2 producing cells, we found that, surprisingly, expression of one or more IRs associated with higher cytokine production (**Fig. 3C-D**). Cells expressing PD-1 and Tim-3 expressed comparable levels of IFN-γ relative to cells singly expressing Tim-3, while cells expressing only PD-1 had significantly reduced IFN-γ production (**Fig. 3C**). In contrast, for IL-2, a significantly higher fraction of cells co-expressing PD-1 and Tim-3 produced IL-2 than cells expressing only one of the receptors (**Fig. 3D**). These data show that expanded Moc22 TILs, which express varying levels of PD-1 and Tim-3, have differential ability to produce cytokines, which may suggest core programming related to activation rather than bona fide exhaustion in this *ex vivo* setting (17, 18).

Prior reports suggest that TILs expressing both PD-1 and Tim-3 are highly dysfunctional in the tumor, and that combination PD-1/Tim-3 blockade promotes their functional activity (12). Therefore, if adoptively transferred into the tumor, where ligands for PD-1 and Tim-3 may be expressed, it is possible that these ACT products would benefit from combination with PD-1 or Tim-3 blockade to promote T cell activation and improved cytokine release in the tumor.

### CD4^+^ and CD8^+^ T cells express distinct combinations of inhibitory receptors in human TILs

Given the high predominance of inhibitory receptors expressed in murine TILs from oral cavity cancer and B16F10 melanoma cell lines, we posited that TILs expanded from human solid tumors would similarly express multiple inhibitory receptors, including PD-1, Tim-3, and Lag-3. While it is known that TILs are often comprised of a heterogeneous T cell population, it is unclear how the expression of IRs including PD-1 compares among CD4^+^ and CD8^+^ TIL populations. Additionally, as the phenotypes of TIL expanded *ex vivo* from human oral cavity cancer are relatively unexplored, we sought to reveal relationships of PD-1, Tim-3, and Lag-3 expression in CD4^+^ and CD8^+^ OCC TILs.

After four weeks of successful TIL expansion from 9 oral cavity tumors (**Fig. 4A**), we profiled the frequency of CD4^+^ and CD8^+^ populations within TIL and determined the frequency of PD-1 expression within each population. We found that the frequency of CD4^+^ and CD8^+^ populations varied depending on the patient (**Fig. 4B**). We hypothesized that more of the CD8^+^ T cells would express PD-1, but interestingly, a greater frequency of CD4^+^ TILs expressed PD-1 (**Fig. 4C**). No difference in Tim-3 between T cell populations was observed in this patient cohort, while Lag-3 was more highly expressed on CD8^+^ T cells (**Fig. 4C**). To understand whether these characteristics were specific for OCC tumors, we expanded TIL from 3 prostate tumors and found that the inhibitory receptor expression by T cell subset was similar to OCC TIL (**Fig. 4B-C**).

**Figure 4:**
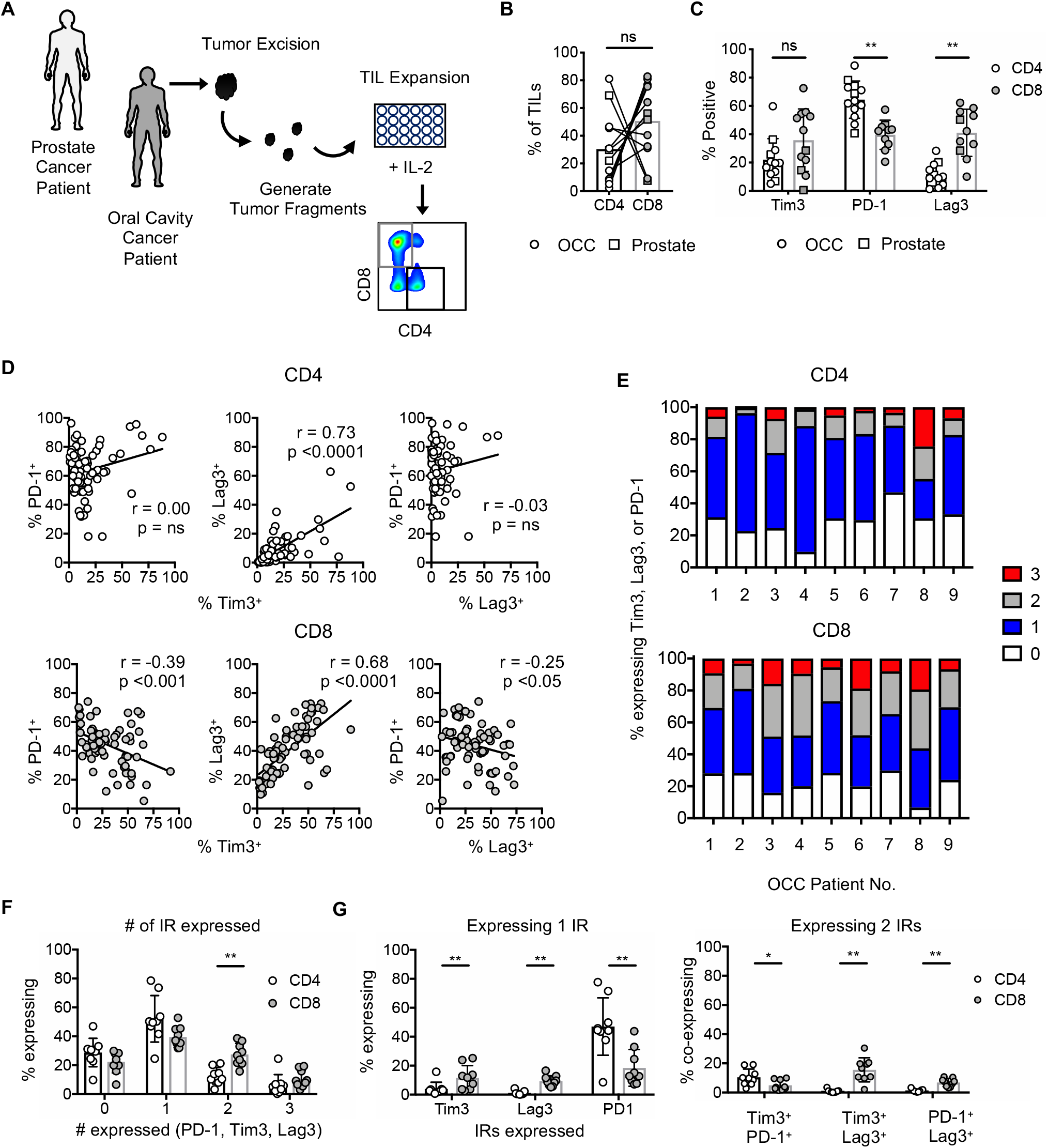
Inhibitory receptor expression dynamics in T cell subsets expanded from human solid tumors. A) TIL expansion schematic. Tumor biopsies were obtained from patients with prostate cancer or oral cavity squamous cell carcinoma (OCC). Tumors were diced and plated in 6000IU IL-2/mL and expanded for 4 weeks prior to analysis. Gating strategy for CD4^+^ and CD8^+^ T cell subsets shown. B) Frequency of CD4 and CD8 T cells in expanded TIL. n=3 prostate cancer, n=9 OCC. C) Frequency of cells expressing inhibitory receptors in prostate and OCC patients. D) Correlation of cells expressing pairs of inhibitory receptors from all TIL cultures of OCC patients. E-G) OCC TILs were assessed for co-expression of Tim-3, PD-1, and Lag-3. Cells can express from 0 to 3 of these receptors. E) Fraction of all TILs within CD4^+^ and CD8^+^ T cell populations expressing 0, 1, 2, or 3 IRs displayed per OCC patient. n=9 OCC patients. F) Statistical comparison of data displayed in E. G) Frequency of T cells expressing only one or two inhibitory receptors, defined and compared between T cell subsets. n=9 OCC patients. Statistical tests: B, C, F, G) Wilcoxon sign rank test, D) Spearman correlation. **p<0.01; ns, not significant.

Moving forward, we sought to better understand PD-1, Tim-3, and Lag-3 expression dynamics within our OCC patient population using all TIL cultures established from 9 patients. Our questions were threefold: 1) does expression of one IR directly correlate with the others, 2) do most of the TILs express more than one of these receptors, and 3) if TILs express 1 or more IRs, which combinations are the most frequent? Understanding of these relationships will be important for design of combination therapies to elicit the most durable antitumor immunity in response to treatment with autologous TIL.

Correlation patterns of PD-1, Tim-3, and Lag-3 pairs were distinct for CD4^+^ and CD8^+^ TILs (**Fig. 4D**). In CD4^+^ TILs, Tim-3 and Lag-3 expression correlated but the frequency of cells involved was relatively low. Additionally, the co-expression of PD-1/Tim-3 or PD-1/Lag-3 did not show a visible trend (**Fig. 4D**). In contrast, for CD8^+^ TILs, PD-1 expression negatively correlated with Tim-3 and Lag-3, while the Lag-3 and Tim-3 pair demonstrated a strong positive correlation (**Fig. 4D**). Overall, these data suggested that TILs expressing PD-1 did not frequently co-express Tim-3 or Lag-3, while TILs expressing Tim-3 were highly likely to co-express Lag-3.

We next questioned how frequent populations of TILs expressing from 1 to 3 inhibitory receptors were in our cohort of OCC patients. For both CD4^+^ and CD8^+^ TILs, over 40% of the cells expressed only one IR, followed in frequency by populations expressing 0, 2, or 3 IRs, suggesting that only a small population concurrently expressed PD-1, Tim-3, and Lag-3 (**Fig. 4E-F**). Within the cohort of cells expressing only one IR, as in **Fig. 4G**, more CD8^+^ T cells singly expressed Tim-3 and Lag-3 relative to CD4^+^ TILs, while the opposite relationship was observed for PD-1. Finally, of TILs expressing 2 IRs, more CD4^+^ TILs co-expressed Tim-3 and PD-1 relative to CD8^+^ TILs, while more CD8^+^ TILs co-expressed Tim-3/Lag-3 and PD-1/Lag-3 relative to CD4^+^ TILs (**Fig. 4G**). Thus, combination therapy targeting these checkpoint molecules would be likely to differentially impact CD4^+^ and CD8^+^ TILs, a concept important for clinical trial design for patients with OCC.

## Discussion

The first evidence that autologous TILs could elicit an antitumor response in patients dates back to 1988 in a cohort of patients with metastatic melanoma(19). Since then, TIL expansion and treatment protocols have incrementally improved to include the incorporation of lymphodepletion preparative regimens, enrichment of tumor-reactive T cells, and addition of costimulatory agents into the rapid T cell expansion process(20, 21). Additionally, there is now robust evidence that TIL therapy is beneficial for patients with solid tumors beyond melanoma, including cholangiocarcinoma (2), cervical (22), colorectal (23), and breast cancers (24). Yet, durable responses to TIL are still only seen in a subset of patients, highlighting the need for novel approaches to prevent relapse (21).

Improvements in TIL efficacy could be licensed by either streamlining the expansion process or by identifying agents which could combine with TIL to prevent T cell exhaustion and extend durability of response. For example, one important question is whether TIL therapy would benefit from combination with checkpoint inhibitors, or whether TIL therapy is even effective in patients after checkpoint blockade (25-27). There is some evidence that co-administration of PD-1 blockade is feasible and safe in TIL recipients (24, 28), but we await clinical trials demonstrating whether the combination can improve responses or survival in larger patient cohorts. As the process of generating TIL on a patient-specific basis is costly and time-consuming, use of preclinical protocols which 1) mirror the clinical process of *ex vivo* TIL expansion, 2) are controlled and reproducible, and 3) could apply across multiple tumor models could help address these questions prior to clinical trial execution.

Herein, we sought to expand our understanding of TIL in malignancies less well studied in the T cell therapy setting. We report successful expansion of TILs *ex vivo* from murine OCSCC and melanoma tumors, while expansion from lung tumor lines yielded few cells. Of the two oral cavity cancer cell lines investigated, CD8^+^ TILs expanded in higher numbers from the Moc22 tumor, which is known to be immunogenic, infiltrated by CD8^+^ T cells, and is responsive to PD-1 blockade (14). In comparison, we obtained fewer TILs from the less immunogenic Moc2, classically unresponsive to PD-1 blockade and with a T cell infiltrate skewed toward CD4^+^ T cells (14). Even so, we were surprised to generate any TIL from the Moc2 tumor as it had been described to have a scant T cell infiltrate (14). Moc22 TILs were phenotypically activated— expressing higher PD-1 and Tim-3 relative to Moc2 TILs—which may indicate that these cells are specific for tumor antigens. In contrast to TILs from the OCC cell lines, TILs from the Lewis lung carcinoma cell line expanded poorly or in many cases not at all. Whether the failure of TIL expansion is related to T cell infiltration, or infiltration of suppressive immune cells remains to be explored. Future work could include manipulation of expansion conditions, including additional cytokine growth factors, or addition of costimulatory agonists or checkpoint inhibitors to promote a more favorable environment for T cell growth. These experiments could be informative for situations where TILs expand poorly or fail to expand from a patient’s tumor.

Mouse TILs overall expressed higher PD-1 and Tim-3 relative to peripheral blood or secondary lymphoid organs. While we did not select for tumor-reactivity within the murine oral cavity TILs, the B16F10 transgenic TIL (specific for gp100 in melanoma) maintained high expression of PD-1 and Tim-3 relative to transgenic cells obtained from the spleen. To improve the clinical relevance of transgenic TIL models, researchers could consider isolating transgenic T cells from tumors rather than lymphoid organs in order to recapitulate phenotypes resembling human TIL. In OCSCC patient TILs, only a small frequency of cells expressed all three PD-1, Tim-3, and Lag-3 receptors. While CD4^+^ TILs more commonly singly expressed PD-1, CD8^+^ TILs more commonly expressed Lag-3^+^ or a combination of 2 markers relative to CD4^+^ T cells. Based on these surface expression profiles, our data further highlight that there may be multiple axes of T cell suppression within the tumor, and that CD4^+^ TILs and CD8^+^ TILs could be suppressed by different mechanisms.

Our work challenges researchers to continue investigating how to make TIL therapy more feasible and effective, and highlights that these goals could be addressed using accessible preclinical models. We assume all cancer types will not have the same routes of immune evasion; therefore, multiple models of ACT beyond melanoma are important for clinical translation. Questions we seek to address in the future include whether tumors relapse after ACT because of 1) T cell exhaustion in the tumor (via Lag-3, Tim-3, PD-1, or others), 2) selective pressure on the tumor, or 3) loss/death of tumor-specific T cells. Overall, our goal is to develop TIL protocols for all types of cancer and determine the settings in which combination therapies can converge to extend survival of cancer patients worldwide.

## Author contributions

HK, DN and CP conceptualized and designed this work. HK, AMRR, MW, AS, RC, CD, MB, GORR, and JH performed experiments and collected data. ML provided clinical samples and oversight. MR provided experimental reagents, guidance and critical feedback. All authors critically read and approved the manuscript prior to submission.

## Acknowledgements

We are thankful to Eric Bartee and Dimitri Arthontoulis for experimental support, as well as Kent Armeson for support on statistical design. We are also grateful for support from Corrine Levingston and Kirsten McDanel at the Flow Cytometry & Cell Sorting Core at the Medical University of South Carolina.

## Supplementary Materials

**Table 1:** Complete list of antibodies

| Target | Clone | Fluorophore | Manufacturer |
| --- | --- | --- | --- |
| αm-Vβ13 | MR12-3 | FITC | BD Biosciences |
|  | MR12-3 | APC | BD Biosciences |
| αm-CD4 | RM4-4 | FITC | BD Biosciences |
|  | GK1.5 | APCCy7 | BD Biosciences |
| αm-CD8 | 53-6.7 | PECy7 | BD Biosciences |
|  | 53-6.7 | PerCpCy5.5 | Biolegend |
|  | 53-6.7 | PE | BD Biosciences |
| αm-PD-1 | 29F.1A12 | PerCpCy5.5 | Biolegend |
|  | 29F.1A12 | APCCy7 | Biolegend |
| αm-Tim-3 | RMT-23 | BV421 | Biolegend |
| αm-IFN-γ | XMG1.2 | PE | BD Biosciences |
| αm-IL-2 | JES6-5H4 | PECy7 | Biolegend |
| αh-CD4 | RPA-T4 | V500 | BD Biosciences |
| αh-CD8 | RPA-T8 | PerCpCy5.5 | Biolegend |
| αh-PD-1 | NAT105 | APCCy7 | Biolegend |
| αh-Tim-3 | F38-2E2 | APC | Biolegend |
| αh-Lag-3 | 11C3C65 | PE | Biolegend |

